# Lamin A/C depletion from myofibers and satellite cells in mice reveals selective muscle pathology

**DOI:** 10.64898/2026.09.03.749246

**Authors:** Qi Jin, Kurenai Tanji, Leroy C. Joseph, Ji-Yeon Shin, Howard J. Worman

## Abstract

Mutations in the lamin A/C gene (*LMNA*), which encodes the nuclear lamina proteins lamin A and lamin C (lamin A/C), have been linked to different human diseases affecting different tissues. Most *LMNA* mutations cause cardiomyopathy and muscular dystrophy, such as autosomal dominant Emery-Dreifuss muscular dystrophy. Recent studies to understand striated muscle laminopathies have taken advantage of *Lmna* conditional knockout mice to examine the effects of lamin A/C depletion in cardiomyocytes and cardiac fibroblasts. However, the role of lamin A/C in skeletal muscle has largely been uncharacterized using conditional knockout mice. We used different mouse lines to deplete lamin A/C from specific cell types in striated muscle. Lamin A/C depletion from fetal myofibers and cardiomyocytes led to no observable phenotype in the skeletal muscles despite leading to dramatic heart dilation and early lethality. Depletion of lamin A/C from both skeletal myofibers and satellite cells was lethal, with the most dramatic myopathic abnormalities observed in the intrinsic muscles of the tongue. The presence of lamin A/C in skeletal muscle satellite cells prevented the development of lethal myopathy when the proteins were deleted only from differentiated myofibers. Overall, our results provide a foundation for understanding the roles of lamin A/C in muscle maintenance and development, including the variable skeletal muscle involvement and much more invariant cardiomyopathy in patients with *LMNA* mutations.

## Introduction

The nuclear lamina is composed of intermediate filament proteins called lamins (1–5). Cryoelectron tomography has demonstrated that in the nuclei of somatic mammalian cells lamins form 3.5-nm diameter fibers different than canonical cytoskeletal intermediate filaments (6). In mammals, three genes encode lamins. *LMNA* encodes the A-type nuclear lamins with lamin A and lamin C (lamin A/C), the two main somatic cell isoforms arising by alternative RNA splicing (7). *LMNB1* encodes lamin B1 and *LMNB2* encodes lamin B2 (8, 9).

Mutations in *LMNA* were first linked to autosomal dominant Emery-Dreifuss muscular dystrophy (EDMD) (10). Subsequently, disorders affecting adipose tissue, bone, peripheral nerves, and those with features of accelerated aging were linked to *LMNA* mutations (11). Dilated cardiomyopathy, usually in conjunction with muscular dystrophy, is the most common of the laminopathies and caused by heterozygous *LMNA* mutations leading to haploinsufficiency, amino acid substitutions, in-frame deletions, or aberrant RNA splicing. The same *LMNA* mutation within a family can result in different distributions of skeletal muscle involvement, but most patients eventually develop dilated cardiomyopathy (10–16).

Pathogenic mechanisms underlying striated muscle laminopathies have been difficult to decipher given the various cellular functions of lamin A/C and the variable cell type specific demands on those functions (11, 17). Among these functions, lamin A/C interact with chromatin to regulate gene expression (18). They interact with integral proteins such as emerin to retain them in the inner nuclear membrane and provide structural stability to the nucleus (19, 20). Notably, mutations in *EMD* encoding emerin cause X-linked EDMD that phenocopies the autosomal dominant type caused by *LMNA* mutations (21). The nuclear membranes are continuous with the endoplasmic reticulum membranes, a site of key metabolic pathways and inflammatory responses (22,23). Lamin A also functions in force transmission between the cytoskeleton and the nucleus by binding to SUN proteins of the LINC complex that spans the nuclear envelope (24, 25).

In mice, germline disruption of *Lmna* resulting in loss of lamin A/C leads to dilated cardiomyopathy, muscular dystrophy, and early lethality (19, 26, 27). Mice with point mutations corresponding to those in human patients also develop dilated cardiomyopathy and muscular dystrophy (28–30). More recently, studies have utilized *Lmna* conditional knockout mice to obtain insights into the requirement of lamin A/C in cardiomyocytes and cardiac fibroblasts (31–37). However, studies on the effects of cell type specific lamin A/C depletion on skeletal muscle have been limited, although indirect effects on bone and neuromuscular junctions with depletion from myofibers have been reported (38, 39). To fill this gap, we have characterized mice with depletion of lamin A/C from skeletal myofibers late in embryogenesis or induced postnatally in myofibers or satellite cells.

## Results

### Lamin A/C depletion from fetal striated myocytes causes lethal cardiomyopathy with minimal skeletal muscle pathology

To determine the effects lamin A/C depletion from cardiomyocytes and skeletal muscle myocytes late in embryonic development, we generated mice hemizygous for a transgene that drives Cre expression from the muscle creatine kinase promoter (MCK-Cre mice) with two floxed *Lmna* alleles. The *Lmna*^fl/fl^ mice have *loxP* sites flanking exon 2 and with germline Cre expression die at postnatal day (P)16-18 (40). The *Mck* promoter is active at approximately embryonic day (E) 17 in developing striated muscle, with expression increasing at and soon after birth (41, 42). In MCK-Cre;*Lmna*^fl/fl^ mice, Cre recombinase expression leads to conditional deletion of *Lmna* from cardiomyocytes and myofibers at approximately E17 (Fig. 1A). We demonstrated lamin A/C depletion from isolated cardiomyocytes by immunofluorescence microscopy (Supplementary Fig. S1A). Similarly, there was a reduction in lamin A/C in skeletal myofibers with near total loss of labeling of the nuclear periphery (Supplementary Fig. S1B). All *Lmna*^fl/fl^ mice and MCK-Cre;*Lmna*^fl/+^ mice survived for a follow-up period of 400 days without any overt signs of illness, whereas MCK-Cre;*Lmna*^fl/fl^ mice had a median survival of 28 days, and all died by P35 (Fig. 1B). Normalized body mass (pup mass divided by litter average) did not differ significantly between genotypes at P14 but was significantly reduced in MCK-Cre;*Lmna*^fl/fl^ at P21 (Fig. 1C). Hematoxylin and eosin (H&E)-stained cross sections of hearts at P21 showed profound left ventricular dilatation and wall thinning in MCK-Cre;*Lmna*^fl/fl^ mice (Fig. 1D).

**Figure 1.**
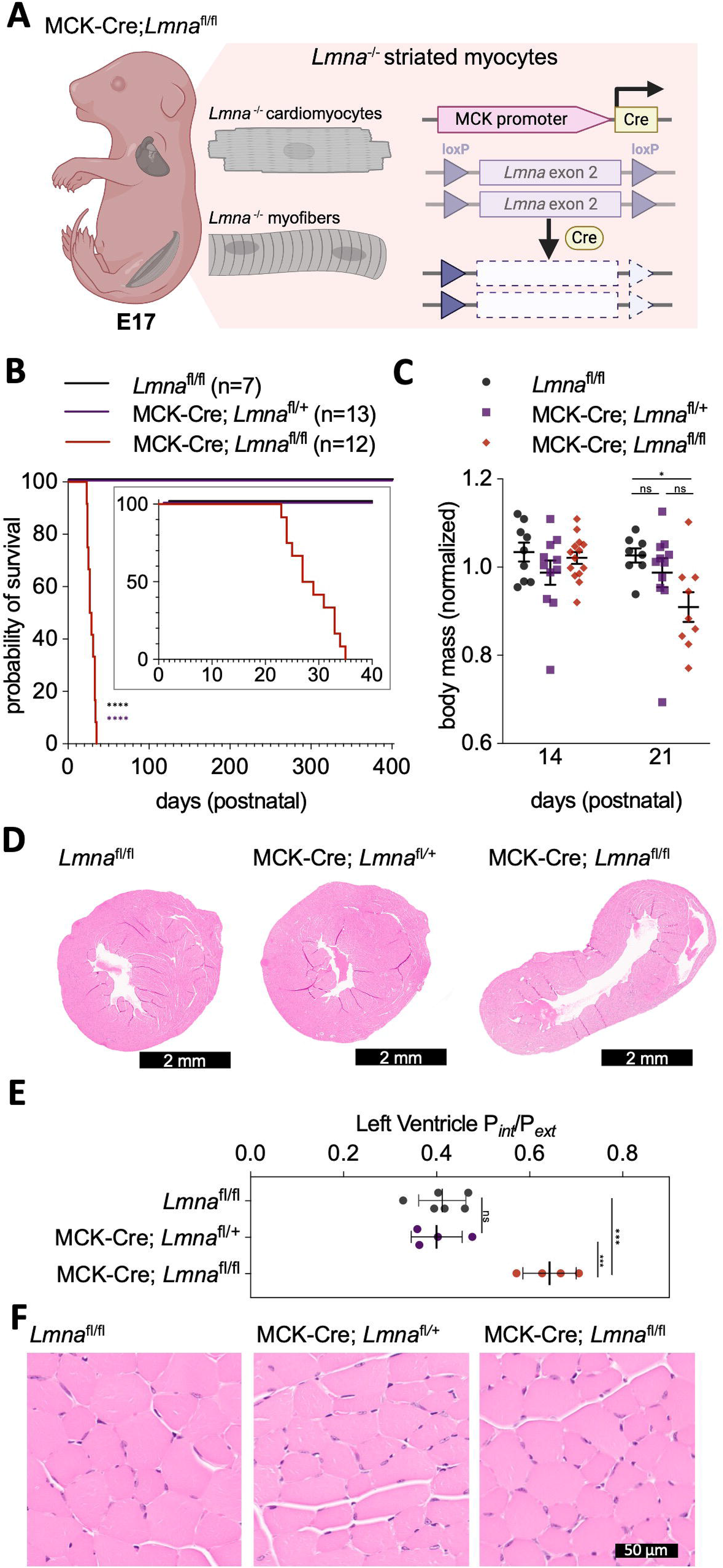
Characterization of mice with depletion of lamin A/C from fetal cardiomyocytes and skeletal myofibers. **(A)** Schematic diagram of MCK-Cre;*Lmna*^fl/fl^ mice. The mice are hemizygous for the MCK-Cre transgene and have *loxP* sites flanking exon 2 of *Lmna*. In cardiomyocytes and skeletal myofibers, Cre excises *Lmna* exon 2 starting at approximately E17 leading to lamin A/C depletion. Created with BioRender.com. **(B)** Kaplan-Meier survival curves for *Lmna*^fl/fl^, MCK-Cre;*Lmna*^fl/+^, and MCK-Cre;*Lmna*^fl/fl^ mice monitored up 400 days postnatal. Inset shows the first 40 days postnatal. MCK-Cre; *Lmna*^fl/fl^ mice had a median survival of 28 days. Statistical significance was evaluated by log-rank (Mantel–Cox) test (χ² = 41.35, df = 2, P < 0.0001). Pairwise log-rank tests with Bonferroni correction confirmed significant shorter survival of MCK-Cre; *Lmna*^fl/fl^ mice compared to *Lmna*^fl/fl^ and MCK-Cre;*Lmna*^fl/+^ mice. Color-coded asterisks next to the survival curve of MCK-Cre;*Lmna*^fl/fl^ mice indicate significance compared to the group corresponding to the color; ****P < 0.0001. **(C)** Normalized body mass at 14 and 21 days of age of *Lmna*^fl/fl^, MCK-Cre; *Lmna*^fl/+^, and MCK-Cre; *Lmna*^fl/fl^ mice. The body mass for each mouse was normalized to the mean body mass of its litter to account for differences in absolute mass of pups between litters. Values are means ± SEM with each colored circle the value for an individual mouse. Normalized body masses were similar between the groups at P14, but significantly different at P21 (One way ANOVA; F = 3.594, P = 0.0425); Šídák’s post-hoc multiple comparisons test found MCK-Cre;*Lmna*^fl/fl^ mice had a significantly lower body mass than *Lmna*^fl/fl^. *P_adj_< 0.05, ns = not significant. **(D)** Representative micrographs of H&E-stained cross sections of hearts at mid-left ventricle level from *Lmna*^fl/fl^, MCK-Cre;*Lmna*^fl/+^ and MCK-Cre;*Lmna*^fl/fl^ mice at P21. **(E)** Ratio of internal perimeter (P_int_) to the external perimeter (P_ext_) of the left ventricle of *Lmna*^fl/fl^, MCK-Cre;*Lmna*^fl/+^, and MCK-Cre;*Lmna*^fl/fl^ mice. Two sections for each biological replicate were measured and means calculated to obtain the ratio for each mouse. Values are means ± SEM with each symbol showing the value for an individual mouse. MCK-Cre;*Lmna*^fl/fl^ mice were significantly different than *Lmna*^fl/fl^ and MCK-Cre;*Lmna*^fl/+^ mice. Statistical significance was determined by one way ANOVA (F = 21.53, P < 0.0001) followed by Šídák’s post-hoc multiple comparisons test. ***P_adj_< 0.0005, ns = not significant. **(F)** Representative micrographs of H&E-stained cross sections of tibialis anterior muscle from *Lmna*^fl/fl^, MCK-Cre;*Lmna*^fl/+^, and MCK-Cre;*Lmna*^fl/fl^ mice at P21.

Trichrome staining showed an increase in fibrosis (Supplementary Fig. S1C). We quantified left ventricular dilation by dividing its inner perimeter by the outer perimeter and the ratio was significantly greater in hearts of MCK-Cre;*Lmna*^fl/fl^ mice (Fig. 1E). In contrast, limb muscle of MCK-Cre;*Lmna*^fl/fl^ mice showed no obvious pathology at P21 (Fig. 1F). The dilated cardiomyopathy that resulted from depletion of lamin A/C from fetal cardiomyocytes was the likely cause of early death in MCK-Cre;*Lmna*^fl/fl^ mice. While lamin A/C was depleted in skeletal myocytes in the same timeframe, it did not result in any histopathological abnormalities in skeletal muscle at the time where the mice began to die. These results show that mice require lamin A/C in cardiomyocytes for early postnatal survival; however, the proteins are not required in skeletal myofibers at the same early ages.

### Survival of mice after depletion of lamin A/C from skeletal myofibers, satellite cells, or both

Induced depletion of lamin A/C from postnatal cardiomyocytes in mice leads to rapid onset cardiomyopathy and death (31, 32, 34–37). We focused on lamin A/C depletion from postnatal skeletal muscle, as such studies have been limited (38, 39). To investigate the effect of lamin A/C depletion in skeletal muscle, we considered not only myofibers but also satellite cells.

Myofibers comprise of most of skeletal muscle volume and are the functional units of force generation (43). Attached to the sarcolemma of myofibers are satellite cells, resident stem cell that can regenerate an entire myofiber (44, 45).

Our approach was to induce cell type specific depletion of lamin A/C in postnatal mice in either myofibers, satellite cells, or both (Fig. 2A). To do so, we generated *Lmna*^fl/fl^ mice with inducible Cre transgenes. For myofiber specific deletion, we used HSA-MerCreMer (HSA-MCM) mice that contain a tamoxifen-inducible human skeletal muscle α-actin gene promoter (46).

**Figure 2.**
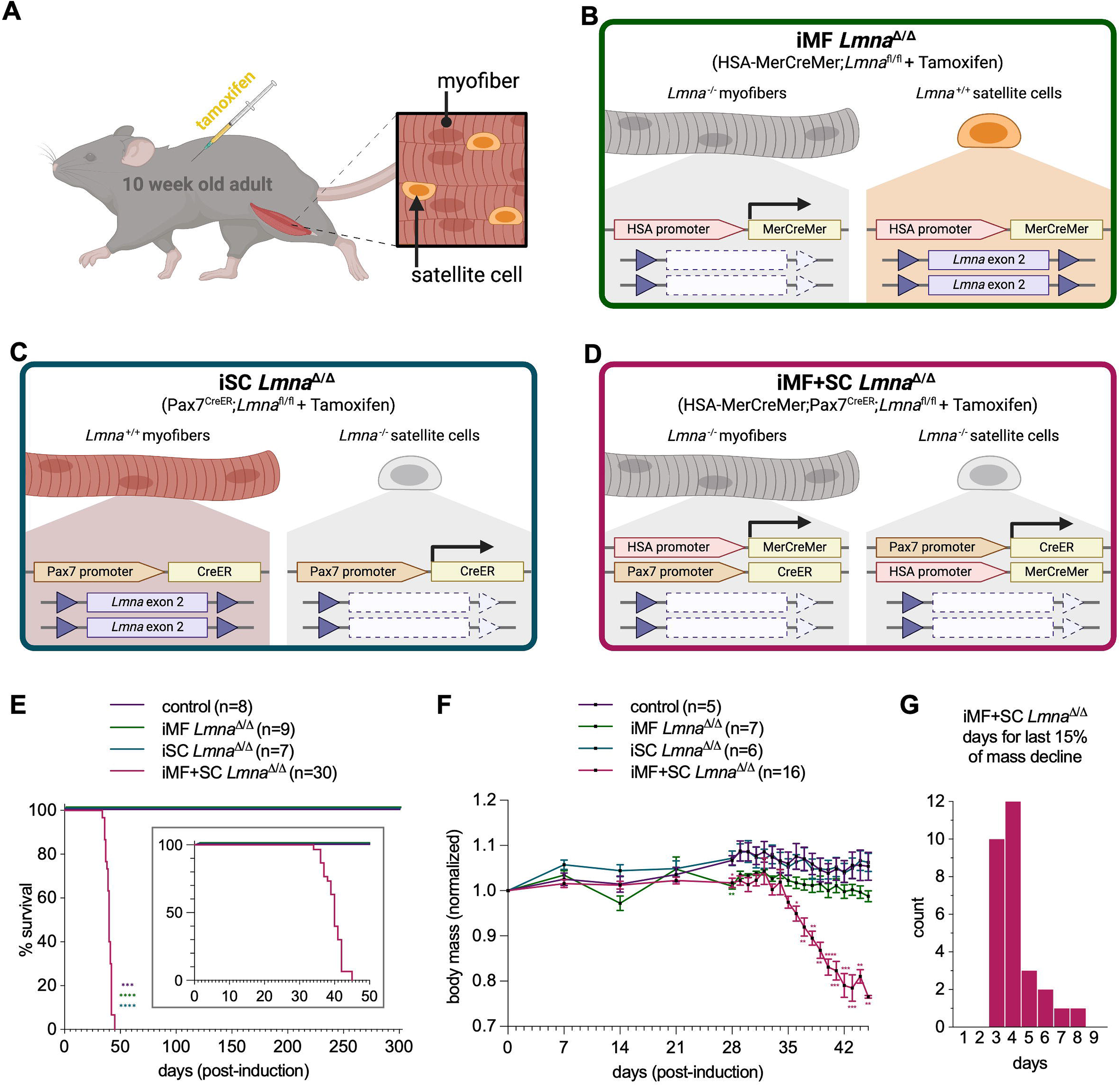
Mouse lines to study lamin A/C depletion from skeletal muscle. **(A)** Schematic diagram showing the experimental design of the study examining depletion of lamin A/C from skeletal muscle cells. At 10 weeks of age, adult *Lmna*^fl/fl^ mice were injected with tamoxifen for induced conditional deletion of lamin A/C from myofibers, satellite cells, or both. (**B**) Schematic diagram of mice with inducible deletion of *Lmna* from mature myofibers (iMF *Lmna*^Δ/Δ^ mice). In mice with genotype HSA-MerCreMer;*Lmna*^fl/fl^, MerCreMer expression is driven by the *HSA* promoter, which is only expressed in mature myofibers and not satellite cells. Injection of tamoxifen results in deletion of exon 2 from both *Lmna* alleles in mature myofibers. **(C)** Schematic diagram of mice with inducible deletion of *Lmna* from satellite cells (iSC *Lmna*^Δ/Δ^ mice). In mice with genotype Pax7^CreER^;*Lmna*^fl/fl^, Cre^ER^ expression is driven by the *Pax7* promoter, which is only expressed in satellite cells and not mature myofibers. Injection of tamoxifen results in deletion of exon 2 from both *Lmna* in satellite cells. **(D)** Schematic diagram of mice with homozygous inducible deletion of *Lmna* from satellite cells (iMF+SC *Lmna*^Δ/Δ^ mice) In mice with genotype HSA;MerCreMer;Pax7^CreER^;*Lmna*^fl/fl^, MerCreMer expression is driven by the *HSA* promoter in mature myofibers, and Cre^ER^ expression is driven by the *Pax7* promoter in satellite cells. Injection of tamoxifen results in deletion of exon 2 from both *Lmna* alleles in both mature myofibers and satellite cells. **(E)** Kaplan-Meier survival curves for control (iMF+SC *Lmna*^Δ/+^), iMF *Lmna*^Δ/Δ^, iSC *Lmna*^Δ/Δ^, and iMF+SC *Lmna*^Δ/Δ^ mice monitored for 300 days post-induction of lamin A/C depletion. Inset shows the first 50 days post-induction. No mortality was observed for control, iMF *Lmna*^Δ/Δ^, and iSC *Lmna*^Δ/Δ^ mice; iMF+SC *Lmna*^Δ/Δ^ mice had a median survival of 40 days. Statistical significance was evaluated by log-rank (Mantel–Cox) test (χ² = 37.08, df = 3, P < 0.0001). Pairwise log-rank tests with Bonferroni correction confirmed significantly shorter survival of iMF+SC *Lmna*^Δ/Δ^ mice compared to control, iMF *Lmna*^Δ/Δ^, and iSC *Lmna*^Δ/Δ^ mice. Color-coded asterisks next to iMF+SC *Lmna*^Δ/Δ^ mouse survival curve indicate significance between itself and the group corresponding to the color; ***P < 0.0005, ****P < 0.0001 **(F)** Normalized body mass of control, iMF *Lmna*^Δ/Δ^, iSC *Lmna*^Δ/Δ^, iMF+SC *Lmna*^Δ/Δ^ mice. Body mass was measured on days 7, 14, 21, 28 and then daily until day 45 post-induction, and the mass of each mouse at each timepoint was normalized to its body mass on the first day of induction. Values are means ± SEM. Statistical significance was evaluated using a restricted maximum likelihood (REML) mixed-effects model with repeated measures (subject matching χ²(1) = 255.9, P < 0.0001; Geisser–Greenhouse ε = 0.2566), confirming significant main effects of group (F(3, 29) = 26.74, P < 0.0001), time (F(5.389, 140.6) = 16.76, P < 0.0001), and time x group interaction (F(16.17, 140.6) = 10.18, P < 0.0001). Color-coded asterisks indicate significant differences, determined by Šídák’s post-hoc multiple comparisons test, between the groups of the corresponding color and control. *P_adj_ < 0.05, **P_adj_ < 0.01, ***P_adj_ < 0.001, ****P_adj_ < 0.0001; unlabeled comparisons are not significant. **(G)** Distribution of iMF+SC *Lmna*^Δ/Δ^ mice (total n = 29) by the number of days to lose the final 15% of body mass before reaching the predetermined humane endpoint. Diagrams in panels A-D were created with BioRender.com.

Administration of tamoxifen to HSA-MCM;*Lmna*^fl/fl^ mice induced homozygous deletion of *Lmna* only the myofibers; we called these animals induced myofiber specific *Lmna*^Δ/Δ^ (iMF *Lmna*^Δ/Δ^) mice (Fig. 2B). We confirmed lamin A/C depletion in myofibers of iMF *Lmna*^Δ/Δ^ mice by immunofluorescence microscopy (Supplementary Fig. S2A). For satellite cell deletion, we used Pax7^CreER^ mice that have a transgene containing an inducible Cre that is controlled by the endogenous *Pax7* promoter (47). In Pax7^CreER^;*Lmna*^fl/fl^ mice, administration of tamoxifen induced homozygous deletion of *Lmna* only in satellite cells; we called these animals induced satellite cell specific *Lmna*^Δ/Δ^ (iSC *Lmna*^Δ/Δ^) mice (Fig. 2C). We confirmed lamin A/C in satellite cells of iSC *Lmna*^Δ/Δ^ mice by immunofluorescence microscopy (Supplementary Fig. S2B). Additionally, we intercrossed HSA-MCM;*Lmna*^fl/fl^ and Pax7^CreER^;*Lmna*^fl/fl^ to generate HSA-MCM;Pax7^CreER^;*Lmna*^fl/fl^ mice, in which administration of tamoxifen induced homozygous deletion of *Lmna* in both myofibers and satellite cells, which we called induced myofiber and satellite cell *Lmna*^Δ/Δ^ (iMF+SC *Lmna*^Δ/Δ^) mice (Fig. 2D). Because *Lmna*^-/+^ mice live for longer than a year (19, 27), we used HSA-MCM;Pax7^CreER^;*Lmna*^fl/+^ mice that were injected with tamoxifen (iMF+SC *Lmna*^Δ/+^ mice) as controls. All mice were given three consecutive daily doses of tamoxifen starting at 10 weeks of age to induce *Lmna* deletion. The iMF *Lmna*^Δ/Δ^, iSC *Lmna*^Δ/Δ^, and control mice survived longer than 300 days whereas iMF+SC *Lmna*^Δ/Δ^ mice had a median survival of 40 days post-induction (Fig. 2E). The endpoint for these mice in all cases was not natural death but rather the predetermined humane endpoint of euthanasia because of >20% body mass decrease. The iMF+SC *Lmna*^Δ/Δ^ mice experienced a sharp decline in body mass starting approximately 35 days after tamoxifen administration, while the body mass of iMF *Lmna*^Δ/Δ^, iSC *Lmna*^Δ/Δ^, and control mice remained stable (Fig. 2F). Consistent with body mass loss, iMF+SC *Lmna*^Δ/Δ^ mice were smaller than control littermates (Supplementary Fig. S2C). We measured the velocity of body mass decline for each individual iMF+SC *Lmna*^Δ/Δ^ mouse represented as the last 15% decrease prior to the euthanasia endpoint (Supplementary Fig. S2D). We observed a distribution of 3 to 8 days heavily skewed towards the lower end, with a median of 4 days for 15% body mass loss (Fig. 2G). These results show mice require lamin A/C in myofibers and satellite cells whereas depletion from each of these cell types individually is compatible with long-term survival.

### Skeletal muscle histopathology in mice with lamin A/C depletion from myofibers, satellite cells, or both

We hypothesized that combined myofiber and satellite cell depletion of lamin A/C would lead to severe histopathological abnormalities in skeletal muscle by the time the animals reached their euthanasia endpoint. To test this hypothesis, we collected skeletal muscle from iMF+SC *Lmna*^Δ/Δ^ mice after euthanasia once they lost >20% body mass, which ranged from 34 to 45 days post-induction of lamin A/C depletion, and from iMF *Lmna*^Δ/Δ^ mice, iSC *Lmna*^Δ/Δ^ mice, and control mice 42 days post-induction.

Microscopic examination of H&E-stained and trichrome-sections sections of quadriceps (rectus femoris) from iMF+SC *Lmna*^Δ/+^ control mice showed normal tissue (Fig. 3A). Quadriceps from most iMF *Lmna*^Δ/Δ^ mice with depletion of lamin A/C from myofibers also appeared normal (Fig. 3B). In some of the replicates of iMF *Lmna*^Δ/Δ^ mice, necrotic myofibers and regenerating myofibers were very sparsely present among the majority of normal myofibers (Supplementary Fig. S3A). Quadriceps sections from iSC *Lmna*^Δ/Δ^ mice with depletion of lamin A/C from satellite cells appeared mostly normal (Fig. 3C). Quadriceps sections from most of iMF+SC *Lmna*^Δ/Δ^ mice with depletion of lamin A/C from both myofibers and satellite cells surprisingly showed no or minimal pathology (Fig. 3D). The minority that did had fibers that exhibited focal necrosis and regeneration (Supplementary Fig. S3B). For a comprehensive analysis, a neuromuscular diagnostic pathologist blind to genotype examined all quadriceps cross sections stained with H&E and trichrome. The pathologist scored severity of atrophy and fibrosis and frequency of necrotic and regenerating fibers of individual samples relative to one another based on expert diagnostic judgment. Scores were then converted to ordinal grades for statistical comparisons (see Materials and methods). Only iMF *Lmna*^Δ/Δ^ mice had a significantly increased severity of skeletal muscle atrophy (Fig. 3E). None of the quadriceps from the four different groups had fibrosis (Fig. 3F). Necrotic fibers were present in 50% of quadriceps of iMF *Lmna*^Δ/Δ^ mice and 17% of iMF+SC *Lmna*^Δ/Δ^ mice but neither was significantly different from control (Fig. 3G).

**Figure 3.**
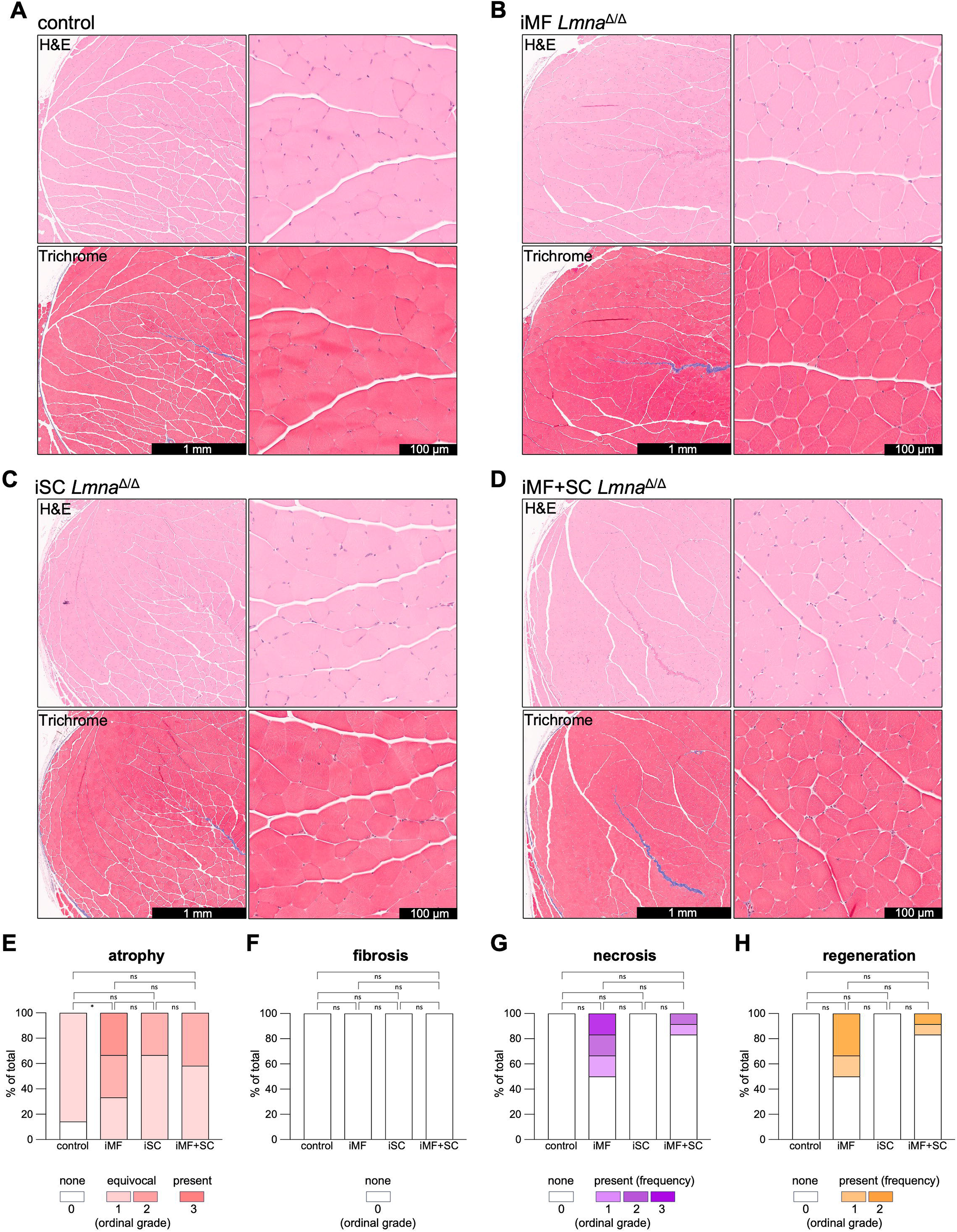
Histopathological assessment of quadriceps muscle. **(A-D)** Representative micrographs of H&E-stained (top) and trichrome-stained (bottom) sections at low (left) and high (right) magnifications of quadriceps of control (iMF+SC *Lmna*^Δ/+^) mice (A), iMF *Lmna*^Δ/Δ^ mice (B), iSC *Lmna*^Δ/Δ^ mice (C), and iMF+SC *Lmna*^Δ/Δ^ mice (D). The internal tendon of the rectus femoris is visible at low magnification. **(E-H)** Histopathological scoring by a blinded pathologist of fiber atrophy (E), fibrosis (F), fiber necrosis (G), and regeneration (H) in quadriceps of control (n = 7), iMF *Lmna*^Δ/Δ^ (iMF; n = 6), iSC *Lmna*^Δ/Δ^ (iSC; n = 6), and iMF+SC *Lmna*^Δ/Δ^ (iMF+SC; n = 12) mice. Stacked bar graphs show the percentage of slides (one per biological replicate) falling into each severity grade, color-coded from least to most severe. For continuous structural changes of atrophy and fibrosis, tissues were scored qualitatively as either none, equivocal, or present. Equivocal and present were further differentiated based on severity where applicable. For focal necrosis and regeneration, tissues were scored based on their relative frequency of occurrence. Scores for all histological features were translated into an ordinal grade as shown in legend, which was used for statistical analysis. Overall differences in ordinal score distributions were evaluated using a non-parametric Kruskal-Wallis test and only atrophy had significant differences (χ² = 8.54, P = 0.0361). Dunn’s post-hoc pairwise comparisons with Holm-Bonferroni correction determined that only iMF *Lmna*^Δ/Δ^ mice were significantly different from control. *P < 0.05, ns = not significant.

Similarly, regenerating fibers were noted 50% of iMF *Lmna*^Δ/Δ^ mice and 17% of iMF+SC *Lmna*^Δ/Δ^ mice, but neither was significantly different from control (Fig. 3H). We additionally checked other muscles and compared to control mice, we did not observe any consistent histopathological abnormalities in diaphragm of iMF+SC *Lmna*^Δ/Δ^ mice beyond what was observed in quadriceps (Supplementary Fig. S3C). The same was true for tibialis anterior (Supplementary Fig. S3D).

Overall, the histopathological abnormalities in the skeletal muscle groups examined in iMF+SC *Lmna*^Δ/Δ^ mice did not appear severe enough to explain the early lethality observed after depletion of lamin A/C from myofibers and satellite cells.

### Tongue muscles show severe histopathological abnormalities in mice with lamin A/C depletion in myofibers and satellite cells

While puzzled by the lack of severe skeletal muscle pathology in the muscles sampled from iMF+SC *Lmna*^Δ/Δ^ mice, we noticed the tongues of some were extended outside the mouth around the time they experienced a rapid body mass decrease (Supplementary Fig. S4A). We therefore performed a histopathological evaluation of tongue muscle by taking cross sections at the intermolar eminence (Supplementary Fig. S4B). This allowed for analysis of the superior and inferior longitudinal myofibers. Tongues of iMF+SC *Lmna*^Δ/+^ control mice appeared normal (Fig. 4A). Tongues from iMF *Lmna*^Δ/Δ^ mice showed some dystrophic features (Fig. 4B). Tongue from the most severely affected iMF *Lmna*^Δ/Δ^ mouse had myofiber atrophy, necrosis, regeneration, and moderate fibrosis (Supplementary Fig. S4C). Tongues from iSC *Lmna*^Δ/Δ^ mice appeared normal (Fig. 4C). Tongues from iMF+SC *Lmna*^Δ/Δ^ exhibited dramatic myofiber deterioration, and notable fibrosis (Fig. 4D). Even a tongue from an iMF+SC *Lmna*^Δ/Δ^ mouse with the most minimal histopathological abnormalities of that group showed focal necrosis and regenerating fibers (Supplementary Fig. S4D). A pathologist blind to experimental groups scored sections of tongues for atrophy, fibrosis, necrosis, and regeneration in the same manner as for quadriceps. Atrophy was present in 50% of tongues of iMF *Lmna*^Δ/Δ^ mice and all iMF+SC *Lmna*^Δ/Δ^ mice and was significantly more severe in the latter (Fig. 4E). Tongue fibrosis was present in all iMF+SC *Lmna*^Δ/Δ^ mice but only 33% of iMF *Lmna*^Δ/Δ^ mice and was significantly more severe in iMF+SC *Lmna*^Δ/Δ^ mice (Fig. 4F). Necrotic fibers were present in tongues of all iMF+SC *Lmna*^Δ/Δ^ mice at a significantly increased frequency compared to all other groups (Fig. 4G). Similarly, regenerating fibers were present in tongues of all iMF+SC *Lmna*^Δ/Δ^ mice at a significantly increased frequency compared to all other groups (Fig. 4H). Prior to any decline in body mass at 28 days post-induction of lamin A/C depletion, histopathological abnormalities were apparent in the tongues of iMF+SC *Lmna*^Δ/Δ^ mice (Supplementary Fig. S4E). The histopathological abnormalities in tongue can explain the rapid body mass loss in iMF+SC *Lmna*^Δ/Δ^ mice starting approximately 35 days post-induction of lamin A/C depletion from both myofibers and satellite cells.

**Figure 4.**
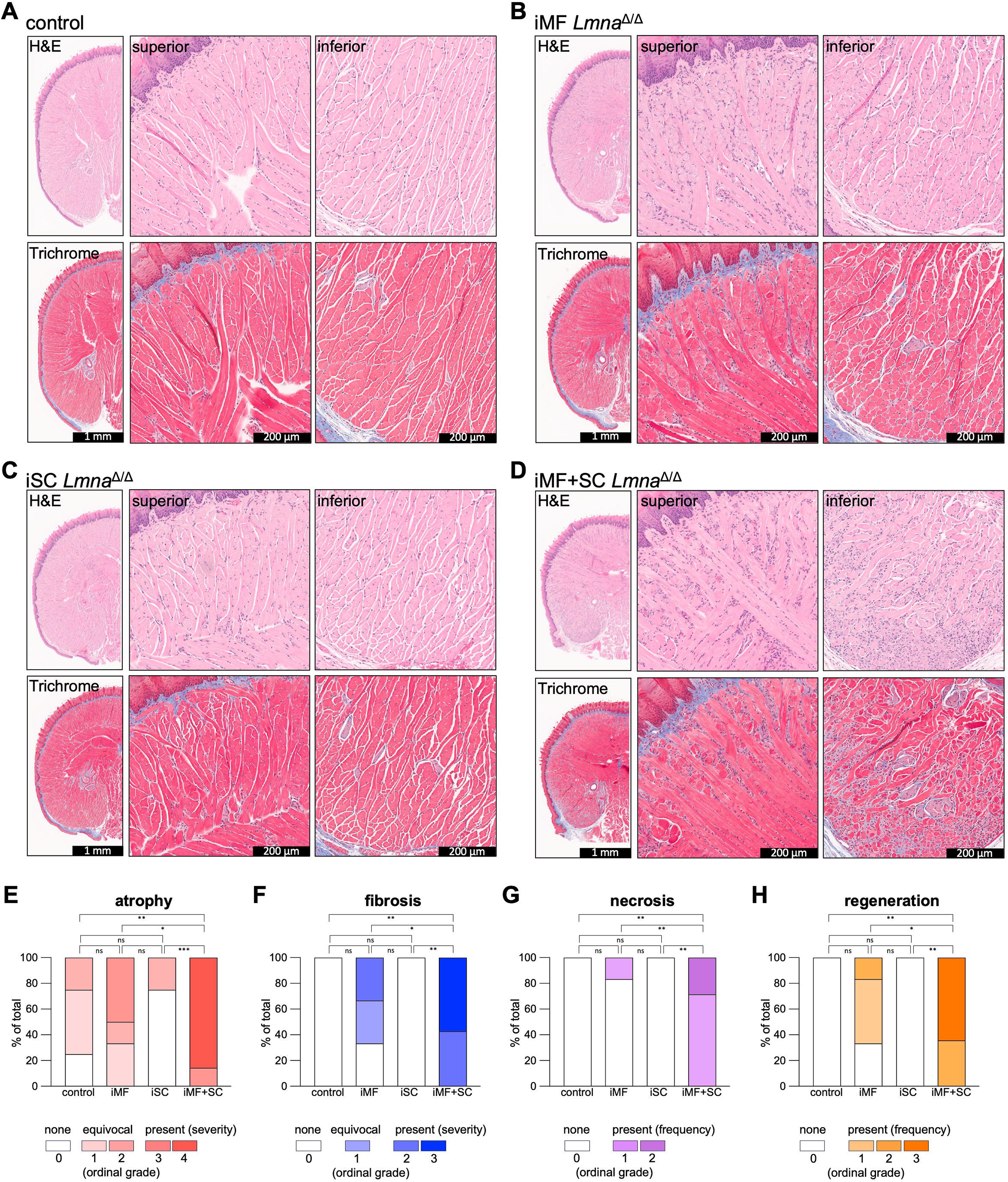
Histopathological assessment of tongue muscle. **(A-D)** Representative micrographs of H&E-stained (top) and trichrome-stained (bottom) sections of tongue of control (iMF+SC *Lmna*^Δ/+^) mice (A), iMF *Lmna*^Δ/Δ^ mice (B), iSC *Lmna*^Δ/Δ^ mice (C), and iMF+SC *Lmna*^Δ/Δ^ mice (D). Higher magnification images to the right show sections of the superior and inferior portions of the tongue as indicated. **(E-H)** Histopathological scoring by a blinded pathologist of fiber atrophy (E), fibrosis (F), necrosis (G), and regeneration (H) in tongues of control (n = 4), iMF *Lmna*^Δ/Δ^ (iMF; n = 6), iSC *Lmna*^Δ/Δ^ (iSC; n = 4), and iMF+SC *Lmna*^Δ/Δ^ (iMF+SC; n = 14) mice. Stacked bar graphs show the percentage of slides (one per biological replicate) falling into each severity grade, color-coded from least to most severe. For continuous structural changes of atrophy and fibrosis, tissues were scored qualitatively as either none, equivocal, or present. Equivocal and present are further differentiated based on severity where applicable. For focal necrosis and regeneration, tissues were scored based on their relative frequency of occurrence. Scores for all histological features were translated into an ordinal grade as shown in legend, which was used for statistical analysis. Overall differences in ordinal grade distributions were evaluated using a non-parametric Kruskal-Wallis test and significant differences were found in atrophy (χ² = 22.7, P = 0.0000459), fibrosis (χ² = 21.5, P = 0.0000846), necrosis (χ² = 22.0, P = 0.0000638), and regeneration (χ² = 22.6, P = 0.0000498). Dunn’s post-hoc pairwise comparisons with Holm-Bonferroni correction determined pairwise significance. *P_adj_ < 0.05, **P_adj_ < 0.01, ***P_adj_ < 0.001, ns = not significant.

### Regeneration of myofibers with lamin A/C depletion

While iMF *Lmna*^Δ/Δ^ mice with depletion of lamin A/C only from myofibers had some histopathological abnormalities in limb and tongue muscles, they survived over 300 days after induction and did not demonstrate significant body mass loss. The presence of lamin A/C in satellite cells appears to have made the difference compared to significant body mass decline of iMF+SC *Lmna*^Δ/Δ^ mice with concurrent depletion of the proteins from satellite cells. When activated, satellite cells regenerate myofibers (44, 45). Pathologists identify active myofiber regeneration on H&E-stained sections by cytoplasmic basophilia and enlarged vesicular nuclei. Internal nuclei are another feature that might also indicate a myofiber has undergone regeneration, although it is not specific marker for regenerated myofibers by itself. In mice, in contrast to humans, internal nuclei persist in myofibers that have fully regenerated (48).

We hypothesized that the iMF *Lmna*^Δ/Δ^ mice did not suffer from severe body mass decline after depletion of lamin A/C from myofibers because of an ability to regenerate damaged ones. To test this hypothesis, we collected quadriceps from iMF *Lmna*^Δ/Δ^ and iMF *Lmna* ^Δ/+^ control mice at 28, 56, 112 days post-induction of lamin A/C depletion and evaluated their H&E stained section for features of regeneration (Fig. 5A). A pathologist blind to experiment group identified a significant higher frequency of myofibers in quadriceps of iMF *Lmna*^Δ/Δ^ mice with histological features of active regeneration at 112 days post-induction (Fig. 5B). We also determined the percentage of myofibers with internal nuclei. At 28 days post-induction both iMF *Lmna*^Δ/Δ^ and iMF *Lmna* ^Δ/+^ control mice had little to no myofibers with internal nuclei, however at 56 and 112 days post-induction iMF *Lmna*^Δ/Δ^ mice had a significantly higher percentage of myofibers with internal nuclei compared to iMF *Lmna* ^Δ/+^ control mice (Fig. 5C). Internal nuclei were also apparent in tongue muscle of iMF *Lmna*^Δ/Δ^ mice 56 days post-induction of lamin A/C depletion (Supplementary Fig. S5). Therefore, depletion of lamin A/C from myofibers led to an increase in muscle regeneration at later timepoints, which likely explained the lack of tongue dysfunction and body mass loss in iMF *Lmna*^Δ/Δ^ mice.

**Figure 5.**
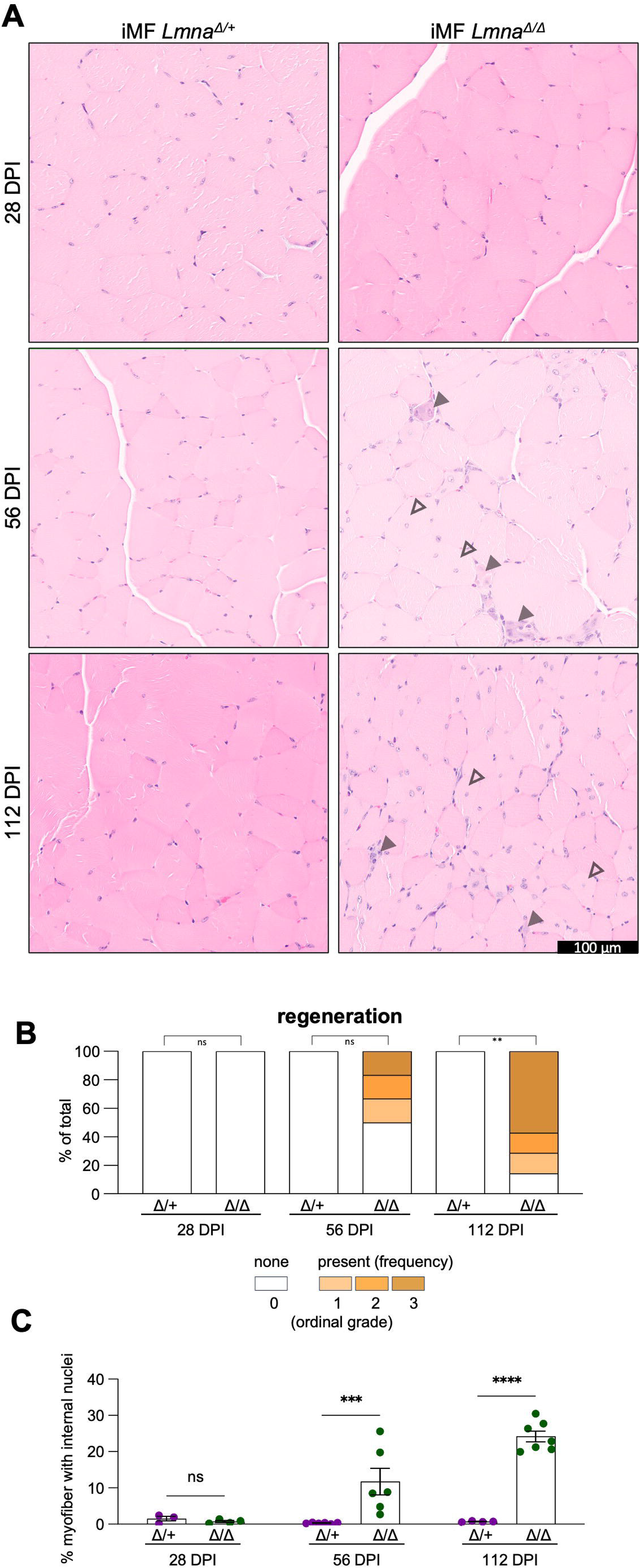
Evidence of myofiber regeneration in iMF *Lmna*^Δ/Δ^ mice. **(A)** Representative photomicrographs of H&E-stained sections of quadriceps of control iMF *Lmna*^Δ/+^ and iMF *Lmna*^Δ/Δ^ mice at 28, 56, and 112 days post-induction (DPI) with tamoxifen to deplete lamin A/C. Solid arrowheads indicate regenerating myofibers and hollow arrowheads myofiber with internal nuclei. (**B)** Histopathological scoring by a blinded pathologist of regenerating fibers in the quadriceps of iMF *Lmna*^Δ/+^ (Δ/+) and iMF *Lmna*^Δ/Δ^ (Δ/Δ) mice. Stacked bar graphs show the percentage of slides (one per biological replicate) falling into each severity grade, color-coded from least to most frequent. n = 3, 6, 4 for iMF *Lmna*^Δ/+^ mice at 28, 56, 112 days post-induction (DPI), respectively; n = 4, 6, 7 for iMF *Lmna*^Δ/Δ^ mice at 28, 56, 112 DPI, respectively. Overall differences in frequency grade distributions were evaluated using a non-parametric Kruskal-Wallis test (χ^2^ = 18.2, P = 0.00267), with time-matched pairwise comparisons evaluated using Dunn’s post-hoc test with Holm-Bonferroni correction. **P_adj_ < 0.01, ns = not significant. **(C)** Percentage of myofibers with central nuclei in iMF *Lmna*^Δ/+^ and iMF *Lmna*^Δ/Δ^ mice at 28, 56, and 112 DPI. Bars represent mean ± SEM; individual data points shown. Two-way ANOVA revealed significant main effects of groups (F(1, 24) = 41.9, P < 0.0001), time (F(2, 24) = 12.9, P = 0.0002), and a significant group x time interaction (F(2, 24) = 14.5, P < 0.0001). Asterisks indicate pairwise Šídák’s post-hoc test comparisons between groups at each time point. *P_adj_ < 0.05, **P_adj_ < 0.01, ****P_adj_ < 0.0001, ns = not significant.

## Discussion

Our results in MCK-Cre;*Lmna*^fl/fl^ mice with depletion of lamin A/C from both fetal cardiomyocytes and skeletal myocytes show that skeletal myofibers are apparently normal up to P21. This highlights the disparity between the requirements for lamin A/C in cardiomyocytes and skeletal myocytes. MCK-Cre;*Lmna*^fl/fl^ mice die from severe cardiomyopathy from P23 to P35. Heart damage and death are consistent findings in postnatal mice early after induced depletion of lamin A/C from cardiomyocytes (32–38). The differential effect of lamin A/C depletion in cardiomyocytes and skeletal myocytes might result from their structural differences and degree of mechanical stress they experience (49, 50). Skeletal muscle is variably affected in humans with *LMNA* mutations, whereas the penetrance of cardiac disease is high (12–16). This further suggests a more critical role for lamin A/C in the heart.

Combined myofiber and satellite cell depletion of lamin A/C in postnatal mice revealed muscle pathology but the effect was selective. At the time of required euthanasia for severe body mass loss, muscles of the tongue were profoundly affected compared to relatively modest pathology in limb muscles and diaphragm. The intrinsic muscles of the tongue have no skeletal attachment to use as leverage for movement or limit its range of motion. Instead, the tongue coordinates the activation of three dimensionally arranged myofibers and utilizes hydrostatic pressure to generate force, change its shape, and move (51). The intrinsic longitudinal myofibers of the tongue, which are largely responsible for protrusion and retraction, were severely affected in iMF+SC *Lmna*^Δ/Δ^ mice. The near-continuous stress of drinking, grooming, and licking in mice require repeated tongue protrusion and retraction. Hence, the myofibers of mouse tongue may be more susceptible to damage from mechanical forces of contraction and stretching in the absence of lamin A/C. Myofibers and satellite cells of the tongue also have a different developmental lineage than trunk and limb muscles (52). The tongue has different regenerative kinetics than limb muscle after acute injury and its satellite cells have different gene expression signatures (53, 54). Although the human tongue similarly functions as a muscular hydrostat (51), it is generally spared in most types of muscular dystrophy. The species difference is likely due to 1) significant differences in the extent of tongue use and 2) the near-complete depletion of lamin A/C from myofibers and satellite cells in iMF+SC *Lmna*^Δ/Δ^ mice versus heterozygous alterations in humans with muscular dystrophy caused by *LMNA* mutations. The severe tongue pathology in iMF+SC *Lmna*^Δ/Δ^ mice with postnatal depletion of lamin A/C does not occur in germline *Lmna*^-/-^ mice prior to death at P42 to P49 (19).

As tongue function is required for eating and drinking, the tongue pathology in iMF+SC *Lmna*^Δ/Δ^ mice was likely responsible for the body mass loss leading to required euthanasia. However, we cannot definitively conclude that tongue dysfunction was the sole cause. While muscles we examined outside the tongue lacked pathology that could explain to the significant body mass decline in iMF+SC *Lmna*^Δ/Δ^ mice, the contribution of other muscle defects cannot be excluded. Myofiber necrosis was present in quadriceps of some iMF+SC *Lmna*^Δ/Δ^ mice at low frequencies and these may have become more abundant if the animals survived longer.

Lamin A/C depletion only from myofibers causes relatively mild pathological alterations, and satellite cells can regenerate them. In the quadriceps of iMF *Lmna*^Δ/Δ^ mice, hallmarks of myofiber regeneration were significantly elevated at 56 days and 112 days post lamin A/C depletion but not at 28 days, indicating the occurrence of muscle damage during this timeframe. The iMF *Lmna*^Δ/Δ^ mice lived and maintained body mass up to at least 300 days after depletion of lamin A/C whereas all iMF+SC *Lmna*^Δ/Δ^ mice suffered from severe body mass loss requiring euthanasia. The key difference is that iMF+SC *Lmna*^Δ/Δ^ mice also had depletion of lamin A/C from satellite cells. The relatively mild pathological changes in iMF *Lmna*^Δ/Δ^ mouse skeletal muscle and their longer survival are similar to what has been reported in mice with constitutive conditional depletion of lamin A/C from myofibers during embryonic development (38, 39). The iSC *Lmna*^Δ/Δ^ mice showed no significant muscle pathology, suggesting that in the absence of myofiber injury, mice do not need lamin A/C in satellite cells. Total ablation of satellite cells from mice shows that they are not required for survival (55). However, skeletal muscle regeneration after acute injury requires functioning satellite cells (56).

It remains to be established if satellite cell dysfunction contributes to the myopathy in autosomal dominant EDMD or related muscular dystrophies caused by *LMNA* mutation. In cultured myoblasts *in vitro*, loss of lamin A/C appears to only reduce the robustness of myogenic differentiation but not block it (57). Skeletal muscle appears to develop normally up until the early postnatal stage in *Lmna*^-/-^ mice, suggesting that there is no primary myogenesis defect in the absence of lamin A/C (19, 27, 58). We cannot exclude the possibility that satellite cells themselves can function normally without lamin A/C and that the proteins are only required after successful differentiation to myofibers. Skeletal muscle of iMF+SC *Lmna*^Δ/Δ^ mice showed evidence of myofiber regeneration. This suggests that satellite cells lacking lamin A/C have some regenerative capacity *in vivo*. However, those fibers do not fully regenerate or deteriorate after full regeneration and are unable to replace damaged myofibers with lamin A/C depletion, resulting in severe and selective muscle pathology.

## Materials and methods

### Animals

The Institutional Animal Care and Use Committee at Columbia University Irving Medical Center approved the protocols. *Lmna*^fl/fl^ (Strain #026284), MCK-Cre (Strain # 006405), HSA-MerCreMer (Strain # 025750), and Pax7^creER^ (Strain # 017763) mice were obtained from The Jackson Laboratory. Mice were housed in a climate-controlled room with a 12-hour light/12-hour dark cycle and fed a standard laboratory diet (Purina Mills, 5053). Both male and female mice were used for experiments as we did not detect any sex-specific differences in survival or pathology. Mice were on a mixed background that was predominantly C57BL/6. See Supplementary materials and methods for details on breeding and follow-up of mice.

### Tissue isolation and processing

Tissues were harvested, quickly washed in PBS, and immediately immersed in 10% formaldehyde (BICCA 3191-1, Neta Scientific). See Supplementary materials and methods for details on processing and staining.

### Histopathological evaluation of skeletal muscle

Histopathological evaluation was performed by a neuromuscular pathologist (K.T.). Slides were assembled by the primary investigator (Q.J.) into study sets. All slides were labeled with only a 3-digit code number, a letter indicating the type of tissue (T for tongue, Q for quadriceps), and the date of slide preparation. Slide labels contained no indication of genotype or experimental group, ensuring the pathologist remained blinded during evaluation. Histopathological scoring was performed using a light microscope, evaluating entire tissue cross-sections. Individual specimens were evaluated comparatively across the blinded cohort based on expert judgment and diagnostic experience. Continuous structural changes of atrophy and fibrosis were evaluated as whole tissue alterations and scored as none, equivocal, or present. Equivocal and present scores were further subdivided on a multi-tier scale based on relative severity when deemed necessary. Focal pathological abnormalities of necrosis and regeneration occurring within individual fibers were scored as none or present, with present further subdivided on a multi-tier scale based on their relative frequency across cross-sections. Following completion of histopathological scoring, the primary investigator converted the scores into ordinal grades with respect to severity or frequency indicated by the pathologist’s evaluation. Samples were then unblinded and the individual grades for each pathological feature were aggregated by experimental group for statistical analysis.

### Counting of myofibers with internal nuclei

5-µm paraffin-embedded sections were obtained from the mid-belly of rectus femoris and stained with H&E. Five randomly selected non-overlapping microscopic fields at 20x magnification were analyzed so that at least 400 myofibers were counted. Internal nuclei were defined as nuclei positioned roughly equidistant from at least two points of the perimeter that spanned at least half of the fiber’s total circumference. Myofibers were classified as having internal nuclei or not; a myofiber with two visible internal nuclei were counted as one.

### Statistics

Overall survival distributions were compared using the log-rank (Mantel–Cox) test with Bonferroni-corrected pairwise comparisons. Longitudinal body mass was evaluated using a restricted maximum likelihood mixed-effects model with Geisser–Greenhouse correction, followed by Šídák’s *post hoc* multiple comparisons test against controls. For other multiple-group comparisons, ordinary one-way or two-way ANOVA followed by appropriate *post hoc* multiple comparisons tests was performed. All aforementioned parametric and survival analyses were performed in GraphPad Prism (v11.0.2). Statistically significant differences in ordinal grade distributions across experimental groups were evaluated using a non-parametric Kruskal-Wallis test followed by *post hoc* pairwise comparisons conducted using Dunn’s test. P-values were adjusted using the Holm-Bonferroni method (reported as P_adj_). Data were analyzed using R (version 4.2.2) with the rstatix and tidyr packages.

## Acknowledgements

We thank the Columbia Medicine Microscopy Core where some of the image processing and analysis was performed.

## Supplementary material

Supplementary material is available online.

## Data availability

All the data used in this study are available from the authors upon request.

## Author contributions

Q. Jin conceived the project, designed methodology, performed experiments, analyzed data, generated figures, and wrote the manuscript; K. Tanji provided blinded histopathological evaluations; L.C. Joseph performed cardiomyocyte isolation; J.Y. Shin conceived the project and designed methodology; H.J. Worman conceived the project, analyzed data, supervised the research, obtained funding, and wrote the manuscript. All authors reviewed the final manuscript.

## Abbreviations

E: Embryonic day
EDMD: Emery-Dreifuss muscular dystrophy
H&E: Hematoxylin and eosin
Lamin: A/C Lamin A and lamin C
P: Postnatal day

