## Supplemental methods, figures, and tables for "Lamin A/C depletion from myofibers and satellite cells in mice reveals selective muscle pathology"

### Supplementary materials and methods

#### Mouse breeding and follow-up

To generate MCK-Cre;*Lmna*$^{fl/+}$ mice, we crossed *Lmna*$^{fl/fl}$ mice with MCK-Cre mice. To generate MCK-Cre;*Lmna*$^{fl/fl}$, we crossed MCK-Cre;*Lmna*$^{fl/+}$ mice *Lmna*$^{fl/fl}$ mice. Mice were weighed at P14 and P21, and weight was normalized to the average of the litter to account for variations in litter size and mother factors that affect the absolute mass of the pups. Daily weighing was not performed because frequent weighing of pups caused earlier lethality despite delicate handling.

We crossed *Lmna*$^{fl/fl}$ mice with HSA-MerCreMer mice to generate HSA-MerCreMer;*Lmna*$^{fl/+}$ mice. To generate HSA-MerCreMer;*Lmna*$^{fl/fl}$ mice, we intercrossed the HSA-MerCreMer;*Lmna*$^{fl/+}$ mice. Similarly, we crossed *Lmna*$^{fl/fl}$ mice to Pax7$^{creER}$ mice to generate Pax7$^{creER}$;*Lmna*$^{fl/+}$ mice, which we intercrossed to generate Pax7$^{creER}$;*Lmna*$^{fl/fl}$ mice. To generate HSA-MCM;Pax7$^{CreER}$;*Lmna*$^{fl/+}$ mice and HSA-MCM;Pax7$^{CreER}$;*Lmna*$^{fl/fl}$ mice, we crossed HSA-MerCreMer; *Lmna*$^{fl/+}$ mice with Pax7$^{CreER}$;*Lmna*$^{fl/+}$. Additionally, we generated Pax7$^{creER/creER}$;*Lmna*$^{fl/fl}$ to cross with HSA-MerCreMer; *Lmna*$^{fl/fl}$ to increase the chance of obtaining HSA-MCM;Pax7$^{CreER}$; *Lmna*$^{fl/fl}$ mice. Genotyping of offspring of all crosses was performed by PCR of DNA isolated from tail clippings using specific primers (Supplementary Table S1).

To generate iMF *Lmna*$^{\Delta/\Delta}$ mice with depletion of lamin A/C from myofibers, we administered 80 mg/kg of tamoxifen by intraperitoneal injections to 10-week-old HSA-MerCreMer;*Lmna*$^{fl/fl}$ mice at a dose of 80 mg/kg/day for three consecutive days. We similarly administered tamoxifen to 10-week Pax7$^{creER}$;*Lmna*$^{fl/fl}$ mice to generate iSC *Lmna*$^{\Delta/\Delta}$ with depletion from satellite cells or 10-week-old iMF+SC *Lmna*$^{\Delta/\Delta}$ mice to generate animals with depletion from both cell types. We also generated iMF+SC *Lmna*$^{\Delta/+}$ and iMF *Lmna*$^{\Delta/+}$ control mice with heterozygous depletion of lamin A/C from myofibers and satellite cells or only myofibers by tamoxifen administration to HSA-MCM;Pax7$^{CreER}$;*Lmna*$^{fl/+}$ mice and HSA-MCM;*Lmna*$^{fl/*}$ mice, respectively.

Mice were followed daily and weighed weekly or more frequently. Mice were euthanized if signs of significant distress were observed, including 1) difficulty with ambulatory movement, 2) failure to eat or drink, 3) >20% decrease in body mass, 4) rough or unkempt coat, and 5) respiratory distress. For experiments examining isolated hearts or skeletal muscle, tissue was isolated immediately after euthanasia. For survival studies, the endpoint was the time or death or when euthanasia was required. Mice were euthanized via isoflurane followed by cervical dislocation as per the approved protocol.

#### Tissue processing and staining

Standard paraffin histological processing and staining were performed at the Columbia University Molecular Pathology Shared Resource. Prior to paraffin mounting, tissues were cut at the site of interest and oriented. Sections of skeletal muscle from the limbs were first cut mid belly, then cross sectional cuts were prepared from the cut site. Sections of tongue were cut at the intermolar eminence. Sections of 5-μm thickness were cut for H&E and Masson's trichrome staining. To obtain frozen sections required for immunofluorescence labeling, tissues were harvested, quickly washed in PBS, then embedded in OCT compound (Leica Biosystems FSC22 CLEAR, Thermo Fisher Scientific). The OCT block was then frozen by submerging in liquid nitrogen chilled isopentane and 10-μm thick sections were then cut.

#### Immunofluorescence microcopy of cardiomyocytes and myofibers

Cardiomyocytes were isolated from mouse hearts using an established protocol (S1). Skeletal myofibers were isolated as described elsewhere (S2). The isolated cells were fixed in formaldehyde and permeabilized with PBS containing 0.5% Triton X-100. Samples were then washed three times with PBS and incubated in blocking buffer (PBS containing 5% bovine serum albumin). Paraffin sections were deparaffinized as described elsewhere (S3). Antigen retrieval was performed by boiling in EDTA buffer (Abcam Antigen Retrieval Buffer, Thermo Fisher Scientific) for 20 minutes. The suspension was then cooled and tissue sections washed with PBS three times. Immunofluorescence labeling of frozen muscle sections was performed as described (S4). Primary and secondary antibodies were used as indicated (Supplementary Table S2). Microscopy was performed on a Zeiss LSM 710 Confocal Microscope (Medicine Microscopy Core at Columbia University Irving Medical Center).

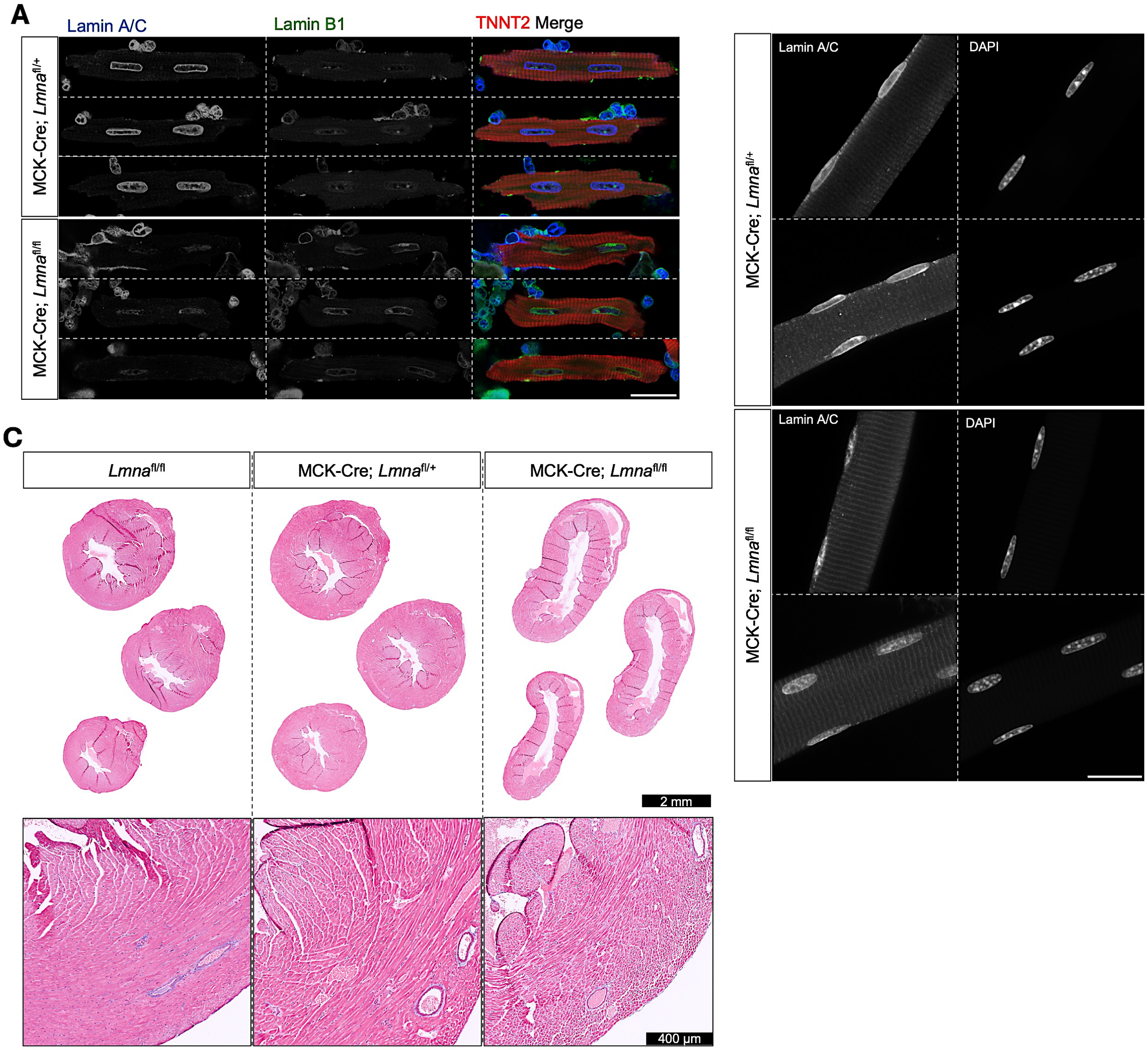


**Supplementary Figure S1.** Confirmation of lamin A/C depletion in cardiomyocytes of MCK-Cre;*Lmna*$^{fl/fl}$ mice and trichrome staining of hearts. **(A)** Representative immunofluorescence micrographs of isolated cardiomyocytes of MCK-Cre;*Lmna*$^{fl/+}$ and MCK-Cre;*Lmna*$^{fl/fl}$ mice labeled with antibodies against lamin A/C, lamin B1, and cardiac troponin T (TNNT2). Three different cardiomyocytes are shown for each genotype. MCK-Cre;*Lmna*$^{fl/fl}$ cardiomyocytes showed near absent labeling while MCK-Cre;*Lmna*$^{fl/+}$ showed prominent nuclear rim staining. Scale bar = 20 µm. **(B)** Representative immunofluorescence micrographs of isolated myofibers of MCK-Cre;*Lmna*$^{fl/+}$ and MCK-Cre;*Lmna*$^{fl/fl}$ mice labeled with antibodies against lamin A/C and stained with 4′,6-diamidino-2-phenylindole (DAPI). Two different myofibers are shown for each genotype. MCK-Cre;*Lmna*$^{fl/fl}$ showed near absent labeling at the nuclear periphery which was prominent in control cells, but with some residual nucleoplasmic fluorescence signal. Scale bar = 20 µm. **(C)** Representative micrographs of trichrome-stained cross sections of left ventricles from *Lmna*$^{fl/fl}$, MCK-Cre;*Lmna*$^{fl/+}$, and MCK-Cre;*Lmna*$^{fl/fl}$ mice at P21. Cross sections were cut at the midpoint of left ventricle (first row), 0.5 mm below (second row), and 0.5 mm below that (third row). The fourth row shows higher magnification micrographs of left ventricle walls.

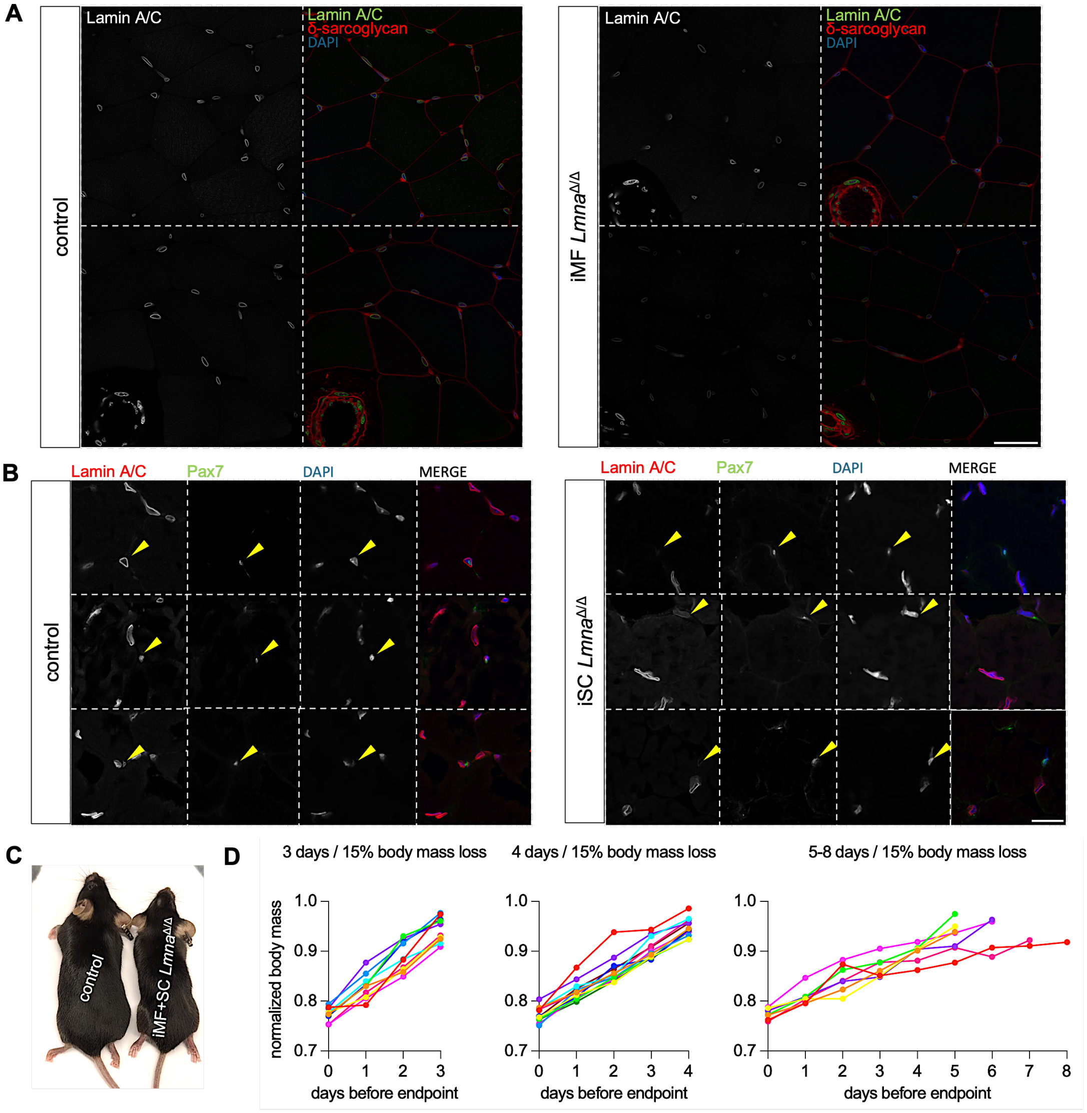


**Supplementary Figure S2.** Tamoxifen-induced depletion of lamin A/C from myofibers or satellite cells in mice. **(A)** Representative immunofluorescence micrographs of quadricep muscle sections from iMF+SC *Lmna*$^{\Delta/+}$ control and iMF *Lmna*$^{\Delta/\Delta}$ mice labeled with antibodies against lamin A/C and δ-sarcoglycan and stained with 4′,6-diamidino-2-phenylindole (DAPI). Two different sections are shown for each genotype. Note blood vessel in lower left corner of each micrograph of iMF *Lmna*$^{\Delta/\Delta}$ mouse muscle, with wall muscle labeled by anti-δ-sarcoglycan antibody and with non-myofiber nuclei expressing lamin A/C. Scale bar = 40 µm. **(B)** Representative immunofluorescence micrographs of quadricep muscle sections from control and iSC *Lmna*$^{\Delta/\Delta}$ mice labeled with antibodies against lamin A/C and Pax7 and stained with DAPI. Arrowheads indicate cells from control mice labeled with both anti-Pax7 antibody and anti-lamin A/C antibody and cells from iSC *Lmna*$^{\Delta/\Delta}$ mice labeled with anti-Pax7 antibody but not labeled with anti-lamin A/C antibody. Three different sections are shown for each genotype. Scale bar = 20 µm. **(C)** Representative photographs of an iMF+SC *Lmna*$^{\Delta/\Delta}$ mouse near required euthanasia endpoint and a littermate control mouse. **(D)** Graphs showing body mass loss trajectories of each individual iMF+SC *Lmna*$^{\Delta/\Delta}$ mouse prior to endpoint of >20% body mass loss. Body mass normalized to body mass at induction is shown as a function of days before the endpoint and divided into 3 plots based on the minimum days it took to lose at least 15% of body mass: 3 days (left, n = 10), 4 days (middle, n = 12), and 5-8 days (right, n = 7). Endpoint (x = 0) is the day the body mass fell below 0.8 normalized body mass (>20% body mass loss). Each colored line represents an individual mouse. Only the ranges of days during which at least 15% of body mass was lost are shown.

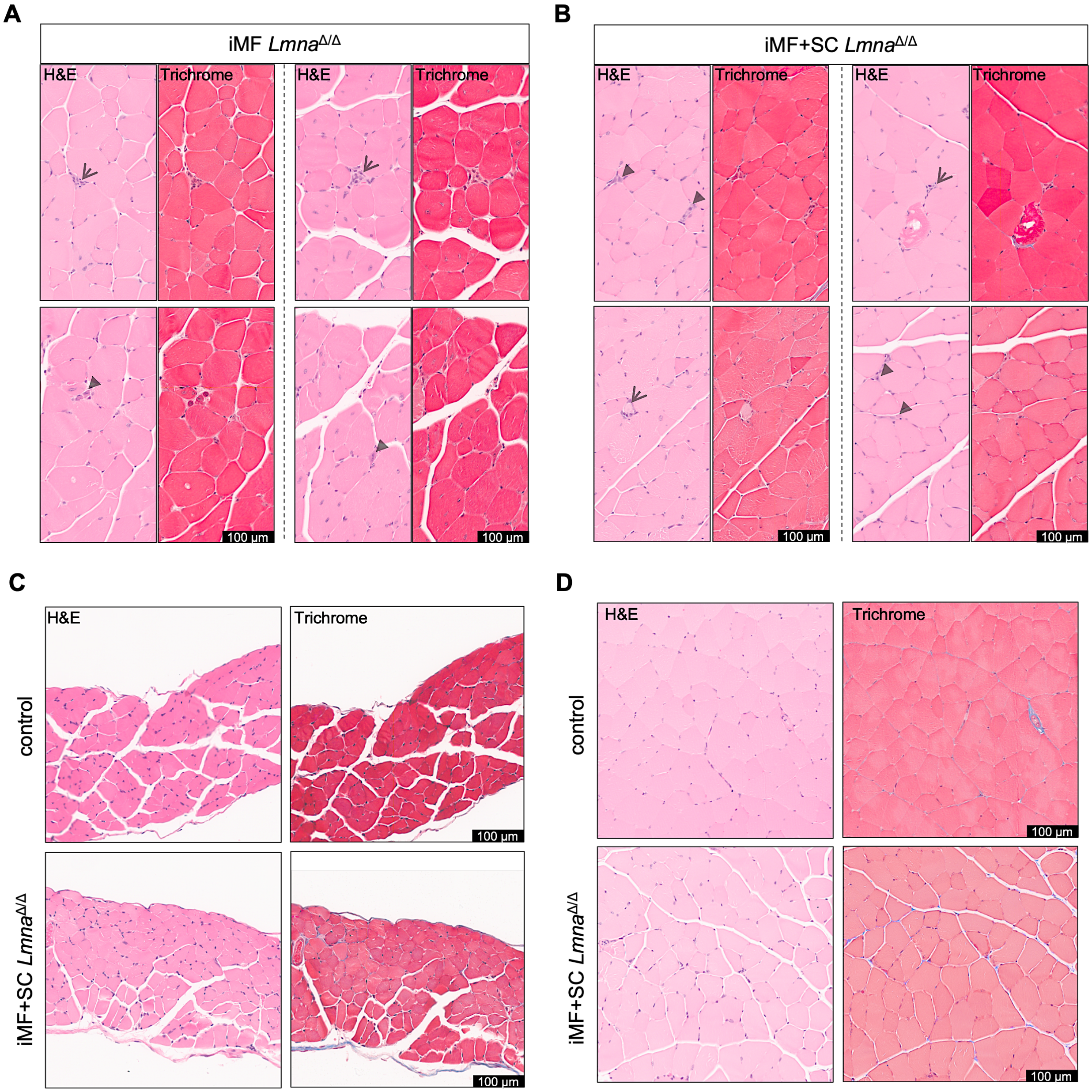


**Supplementary Figure S3.** Histopathological features of skeletal muscle in mice with depletion of lamin A/C from myofibers or both myofibers and satellite cells. **(A)** Representative micrographs of H&E-stained and trichome-stained sections of quadriceps from two biological iMF *Lmna*$^{\Delta/\Delta}$ mouse replicates. Two views of each were selected to specifically show examples of necrosis and regenerating fibers. Stick arrows point to necrotic myofibers, and solid arrowheads point to regenerating myofibers. **(B)** Representative micrographs of H&E-stained and trichome-stained sections of quadriceps from two biological iMF+SC *Lmna*$^{\Delta/\Delta}$ mouse replicates. Two views of each were selected to specifically show examples of necrosis and regenerating fibers. Arrows point to necrotic myofibers and arrowheads point to regenerating myofibers. **(C)** Representative micrographs of H&E-stained and trichome-stained sections of diaphragm from iMF+SC *Lmna*$^{\Delta/+}$ control and iMF+SC *Lmna*$^{\Delta/\Delta}$ mice. **(D)** Representative micrographs of H&E-stained and trichome-stained sections of tibialis anterior from iMF+SC *Lmna*$^{\Delta/+}$ control and iMF+SC *Lmna*$^{\Delta/\Delta}$ mice.

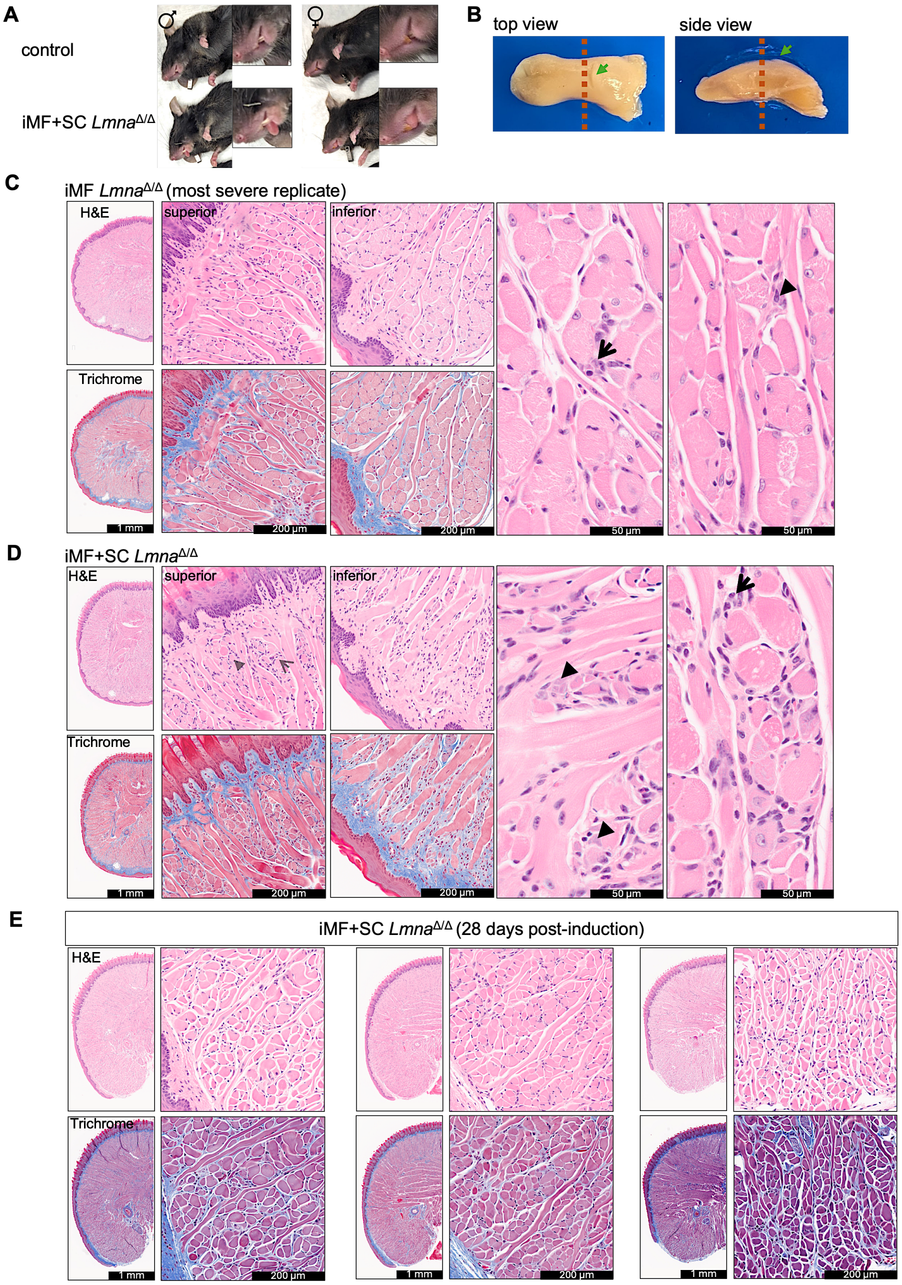


**Supplementary Figure S4.** Histopathological features of tongue muscle in mice with depletion of lamin A/C from myofibers or both myofibers and satellite cells. **(A)** Representative photographs of a sibling pair of male and female iMF+SC $Lmna^{\Delta/+}$ control mice and iMF+SC $Lmna^{\Delta/\Delta}$ mice showing passive extension of the tongues in the iMF+SC $Lmna^{\Delta/\Delta}$ mice. **(B)** Representative photographs of dissected and formaldehyde fixed tongue viewed from the top and side. Dashed lines show where the cross-sectional cuts were made. Arrow points to the intermolar eminence, the marker for the cut. **(C)** Representative micrographs of H&E-stained and trichome-stained sections of tongue from the iMF $Lmna^{\Delta/\Delta}$ mouse that had the most serve overall histopathological abnormalities among animals of that genotype. Stick arrows point to necrotic myofibers and solid arrowheads point to regenerating myofibers. **(D)** Representative micrographs of H&E-stained and trichome-stained sections of tongue from an iMF+SC $Lmna^{\Delta/\Delta}$ mouse. Arrows point to necrotic myofibers and arrowheads point to regenerating myofibers. **(E )** Representative H&E-stained and trichrome-stained sections of tongues from three iMF+SC $Lmna^{\Delta/\Delta}$ mouse replicates collected at 28 days post-induction of lamin A/C depletion.

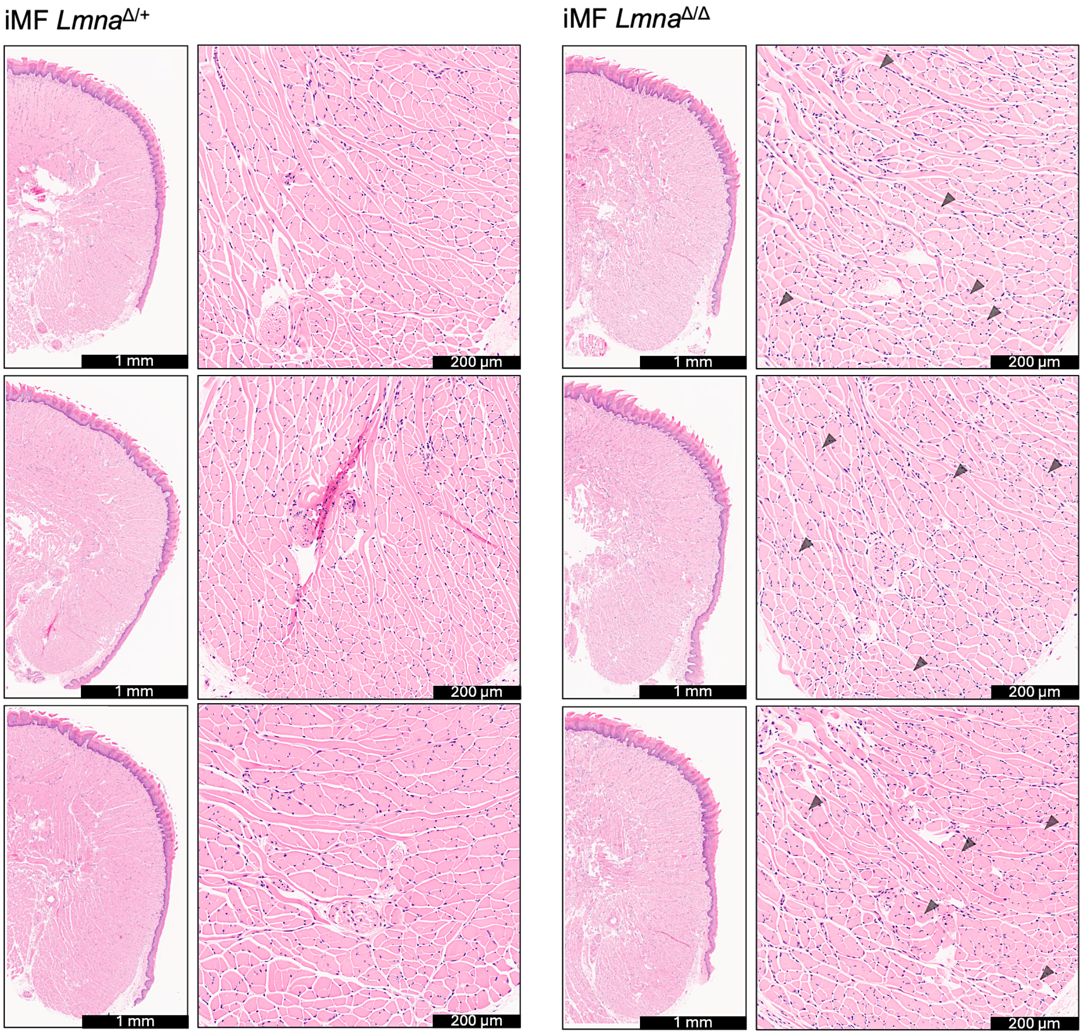


**Supplementary Figure S5.** H&E-stained sections of tongues of iMF *Lmna*$^{\Delta/+}$ and iMF *Lmna*$^{\Delta/\Delta}$ mice at 56 DPI. Arrowheads indicate select representative internal nuclei in higher magnification micrographs of tongue from the iMF *Lmna*$^{\Delta/\Delta}$ mouse.

**Supplementary Table S1.** Sequences of primers for mouse genotyping

| Allele/Transgene | Forward | Reverse |
| --- | --- | --- |
| *Lmna*$^{fl/fl}$ | 5′-AACCCAGCCTCAGAAACTGGTGGATG-3′ | 5′-GACAGCTCTCCTCTGAAGTGCTTGGA-3′ |
| MCK-Cre | 5’-GTGAAACAGCATTGCTGTCACT-3’ | 5’-TAAGTCTGAACCCGGTCTGC-3’ |
| HSA-MCM | 5’-CAGGTAGGGCAGGAGTTGG -3’ | 5’-TTTGCCCCCTCCATATAACA -3’ |
| Pax7$^{CreER}$ | 5′- CTGTGCTGGGACTTCTTCCT -3′ (common) | 5′-AAAGACGGCAATATGGTGGA-3′ |
| Pax7$^{wt}$ | | 5′-AGACTCAGGGCTTGGGAAGG-3′ |

**Supplementary Table S2.** Antibodies for immunofluorescence labeling

**Primary antibodies**

| **Antigen** | **Host Species** | **Source** | **Dilution** |
|---|---|---|---|
| Lamin A/C | Mouse | Santa Cruz [sc-376248] | 1:50 |
| Lamin A/C | Rabbit | Abcam [ab26300] | 1:100 |
| Lamin B1 | Rabbit | (S5)[1] | 1:100 |
| TNNT2 | Goat | Abcam [ab56357] | 1:100 |
| δ-sarcoglycan | Mouse | Vector Laboratories [VP-D501] | 1:100 |
| Pax 7 | Mouse | DSHB [AB_528428] | 1:50 |

**Secondary antibodies**

| **Antigen/Conjugate** | **Host Species** | **Source** | **Dilution** |
|---|---|---|---|
| Mouse IgG/Alexa Fluor™ 488 | Donkey | Invitrogen[2] (A21202) | 1:500 |
| Goat IgG/Alexa Fluor™ 568 | Donkey | Invitrogen (A11057) | 1:500 |
| Rabbit IgG/Alexa Fluor™ 647 | Donkey | Invitrogen (A31573) | 1:500 |
| Rabbit IgG/Alexa Fluor™ Plus 488 | Donkey | Invitrogen (A32790) | 1:500 |
| Mouse IgG/Alexa Fluor™ Plus 647 | Donkey | Invitrogen (A32787) | 1:500 |

[1]See Supplementary material reference S5.

[2]Invitrogen is a Thermo Fisher Scientific brand.

**Supplementary material references**

S1. Joseph, L.C., Kokkinaki, D., Valenti, M.C., Kim, G.J., Barca, E., Tomar, D., Hoffman, N.E., Subramanyam, P., Colecraft, H.M., Hirano, M., et al. (2017) Inhibition of NADPH oxidase 2 (NOX2) prevents sepsis-induced cardiomyopathy by improving calcium handling and mitochondrial function. *JCI Insight*, **2,** e94248.

S2. Pasut, A., Jones, A.E. and Rudnicki, M.A. (2013) Isolation and culture of individual myofibers and their satellite cells from adult skeletal muscle. *J. Vis. Exp.*, **22,** e50074.

S3. Patel, P.G., Selvarajah, S., Boursalie, S., How, N.E., Ejdelman, J., Guerard, K.P., Bartlett, J.M., Lapointe, J., Park, P.C., Okello, J.B., et al. (2016) Preparation of formalin-fixed paraffin-embedded tissue cores for both RNA and DNA extraction. *J. Vis. Exp.*, **114,** 54299.

S4. Hung, M., Lo, H.F., Beckmann, A.G., Demircioglu, D., Damle, G., Hasson, D., Radice, G.L. and Krauss R.S. (2024) Cadherin-dependent adhesion is required for muscle stem cell niche anchorage and maintenance. *Development*, **151,** dev202387.

S5. Cance, W.G., Chaudhary, N., Worman, H.J., Blobel, G. and Cordon-Cardo, C. (1992) Expression of the nuclear lamins in normal and neoplastic human tissues. *J. Exp. Clin. Cancer Res.*, **11**, 233-246.
